# Do orexin/hypocretin neurons track sensory, cognitive, or motor information?

**DOI:** 10.1101/2022.04.13.488195

**Authors:** Eva Bracey, Aditi Aravind, Nikola Grujic, Daria Peleg-Raibstein, Denis Burdakov

## Abstract

Activation of hypothalamic hypocretin/orexin neurons (HONs) is a neural substrate of arousal. HONs activate during sensory stimuli, and are thus thought to regulate arousal according to sensory input. Here, we measured body movements occurring during sound cues or associated reward outcomes, and used an encoding model to ask whether HONs indeed specialize in tracking certain features, or multiplex diverse types of features. Although some single HONs multiplexed feature combinations, during the cue period the overall HON signal primarily tracked body movements. This persisted across cues signaling different reward probabilities, and substantially diverged from reward-probability tracking in concurrently-recorded VTA dopamine neurons. In contrast, during reward outcome, HONs predominantly signaled the presence or absence of reward, and not body movements, nor surprise or reward prediction error. These results describe an unexpectedly specialized and flexible logic of HON activation, suggesting a role for HONs in tracking actions and subsequent reinforcements.

## INTRODUCTION

Arousal plays a key role in cognition, including relationships between actions and reinforcements which are of enduring interest across neuroscience, psychology, economics, and artificial intelligence ^1–19^. A neural substrate of arousal in mammals is the activity of hypocretin/orexin neurons (HONs) of the lateral hypothalamus (LH) ^20–39^. Much research seeking a logic for how the brain allocates arousal thus focused on defining behavioral and sensory features tracked by HONs. These studies produced a growing awareness that HONs respond not only to internal energy state ^27, 40–43^ and to external stimuli across the sensory spectrum^20, 44–47^, but also to other features such as behavioral state ^20, 21, 47–49^, as well as reward-predicting cues^50, 51^. Since such features are often present simultaneously in nature, this raises a fundamental question of whether HONs track particular features, or whether they respond non-selectively to many features. Addressing this question requires a multivariate analysis of multiple co-occurring features and the corresponding neural output ^52–57^. This multivariate approach is important, since it avoids potential misidentification of neural tracking of sensory vs motor features ^52, 58^.

The responses to reward-predicting cues are of particular interest for disentangling neural correlates of associative perception, such as reward prediction ^1, 59, 60^. In Pavlovian conditioning studies typically used to probe such correlates (Fig. 1A), central questions are whether neurons distinguish cues that indicate distinct reward probabilities, and whether neural responses to ensuing reward outcome code ‘reward prediction error’^1, 59, 60^. These questions have been traditionally explored by examining simple bivariate relations between neural activity and reward probability ^61, 62^ or uncertainty ^60^, without taking into account spontaneous motor actions that are increasingly recognized to affect brain-wide neural coding ^52, 55–57, 60^. Such actions can coincide with reward probability cues ^60^, again emphasizing the need for multivariate analysis to address the fundamental question of whether neural responses are explained by the reward or motor features ^52, 54^.

**Figure 1.**
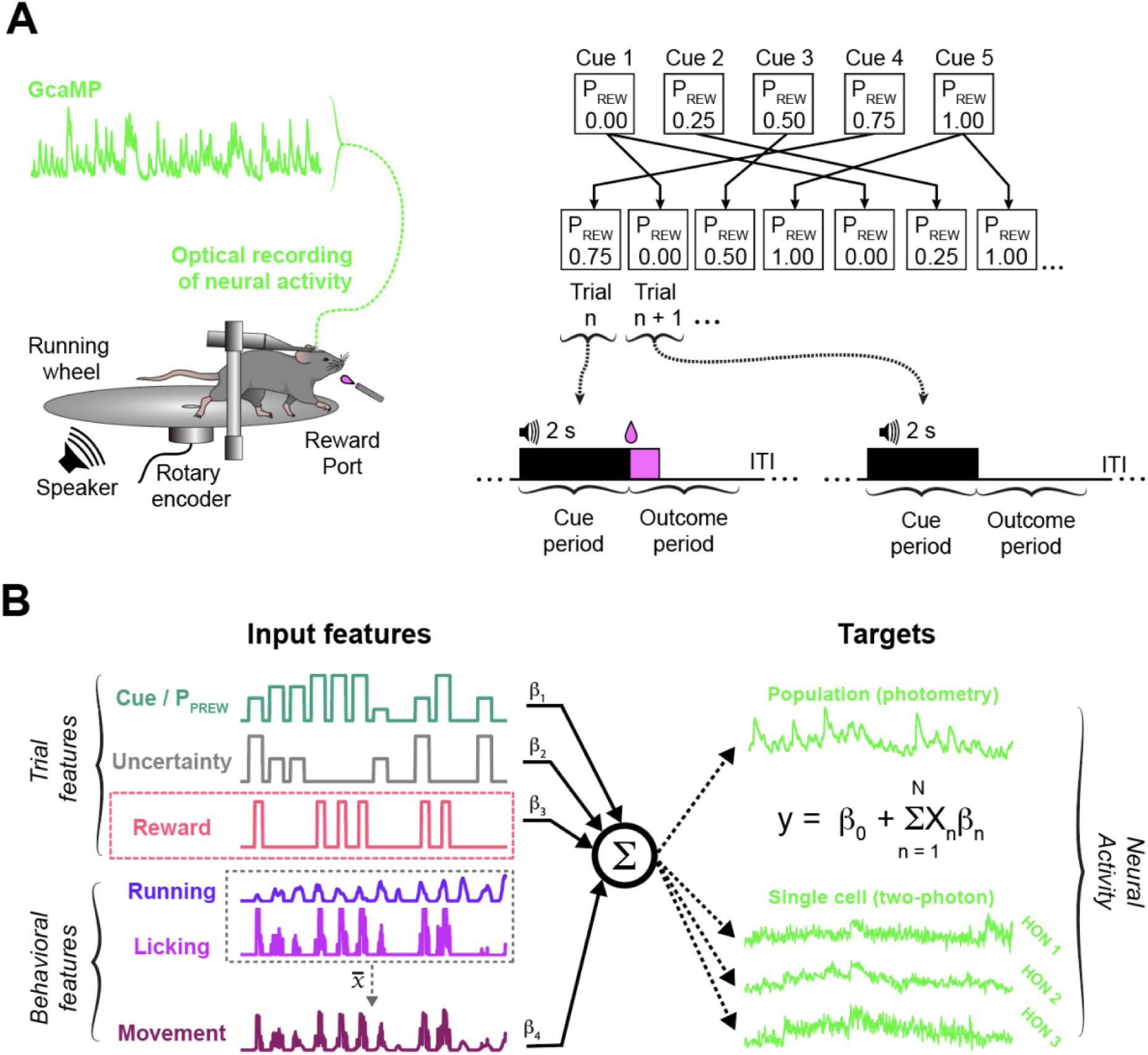
Experimental and analytical approaches used in this study. A. The Pavlovian paradigm. Top, cues were different tones (4 - 12kHz, see Methods) each paired with a distinct reward probability (P_rew_, 0, 0.25, 0.5, 0.75 or 1) in a randomized trial sequence. Bottom, trial structure, illustrating ‘cue period’ (black bar) and ‘outcome period’ (pink bar) analyzed in the study. B. The experimental setup for photometry or 2-photon experiments. C. Multivariate encoding model used to quantify the relationship between trial and behavioral features and neural activity (see Methods).

HON activation logic has not been studied across the full range of reward probabilities, while also considering behavioral state. Therefore, here we measured HON activity during cue and outcome periods of a Pavlovian paradigm with a probabilistic cue-outcome pairing, and concurrent behavioral tracking (Fig. 1A). We analyzed the results first using simple correlations, to relate to classic work using such analyses^60^, and then using a multivariate encoding model ^54^ (Fig. 1B, Fig. S1). With these approaches, we probed the open questions of whether HONs respond to reward probability and/or reward prediction error, and how cue, reward, and motor information is organized across HONs.

## RESULTS

### During the cue period, HONs primarily track body movements

We initially assessed HON activity by fiber photometry of HON-targeted calcium sensors (Fig. 2A, Fig. S2A, see Methods), which is widely-used for assessing bulk HON activity ^37, 45, 46, 63, 64^. In the same mice, we performed photometry recordings from dopamine neurons (DANs) in the ventral tegmental area (VTA) (Fig. 2A-D). The DAN recordings here served a parallel “positive control”, testing whether our experimental paradigm could detect neural coding of reward probability by checking for its known monotonic encoding in DANs (Fig. 2D, compare with ^60^). In trained mice (Fig. S2B), the HON and DAN population responses to cues were not correlated with each other, suggesting that each transmits distinct information (Fig. 2E). Simple analysis (based on^60^) indicated that DAN responses to reward-predicting cues monotonically increased with reward probability as expected ^60^, while HON responses were significantly influenced by uncertainty (Fig. 2D; analysis details are in the figure legends). However, multivariate analysis and predictor assessment (Fig. 1B, see Methods) of the same data did not support this conclusion about HONs, instead indicating that the HON population primarily tracks behavioural state rather than reward-expectation features (Fig. 2F,G). Overall, these results suggest that, during reward-predicting cues, the HON population is not specialized to signal reward expectation features as previously reported for DANs ^60^, but instead primarily tracks body movements.

**Figure 2.**
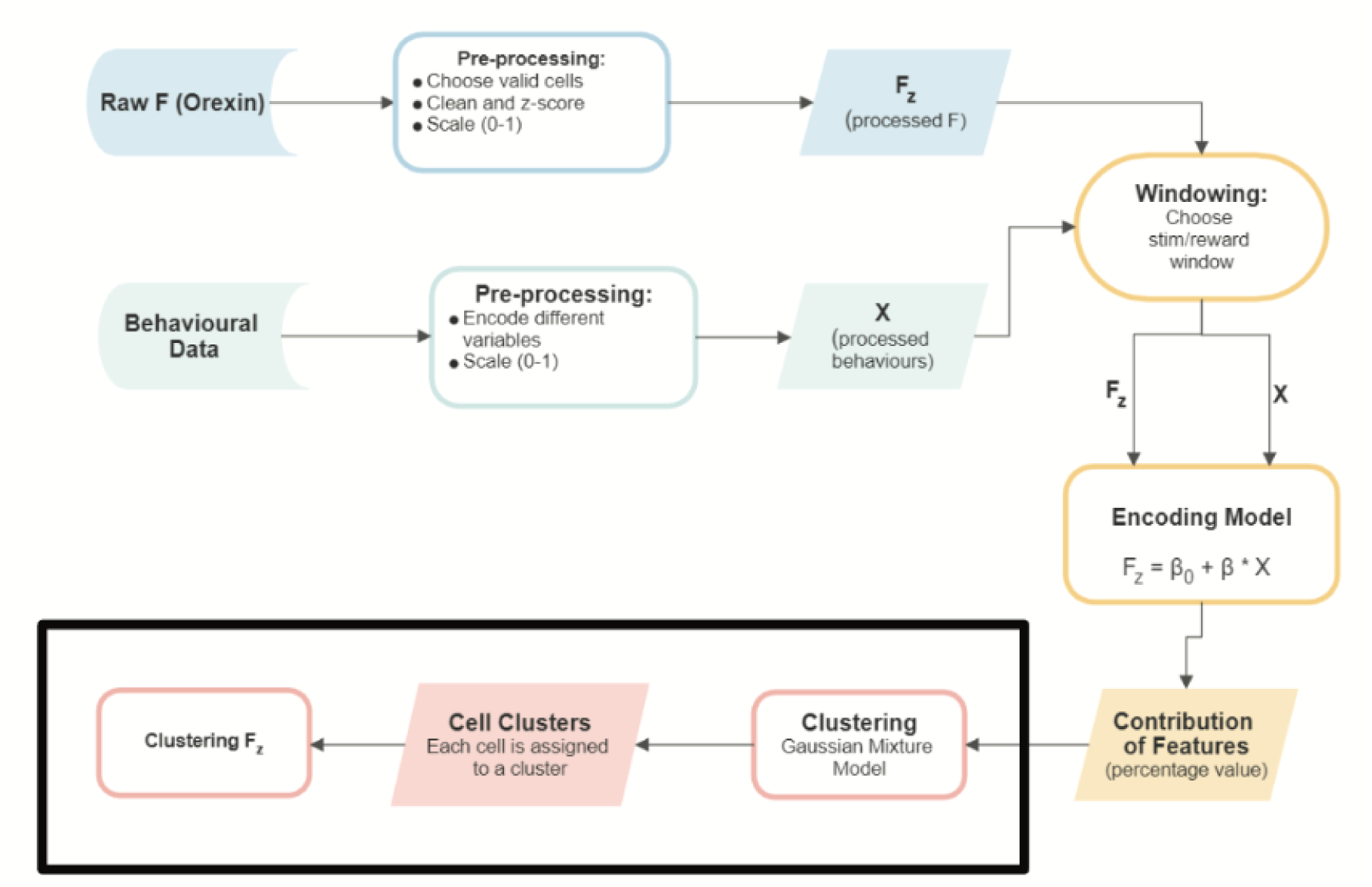
During cue period, HON population primarily tracks body movements. A. Left, coronal brain section showing position of optical fiber and orx-GCAMP expression in the LH. Middle, sagittal section showing viral injection sites of orx-GCaMP in LH (blue), and flex-GCaMP (red) in the VTA. Right, coronal brain section showing position of optical fiber and flex-GCAMP expression in dopaminergic neurons (DANs) of the VTA. *Int: internal capsule, mtt: mammillothalamic tract, ot: optic tract, V3: third ventricle, MM: medial mamillary nucleus, PAG: peri-aqueductal gray, SNR: substantia nigra pars reticularis*. B. Recording setup schematic. C. Responses of LH HON-GCAMP signals (top row, blue) and VTA DAN-GCaMP signals (bottom row, red) to cues signaling the five reward probabilities (P_REW_). n = 10 mice, data are mean +/-SEM. D. Summary of data in C for maximal responses. The same statistical tests used to assess coding of probability and uncertainty in the classic work^60^ indicated that: DAN responses monotonically increase with probability (Spearman r = 1, P < 0.01); HON responses do not monotonically follow probability (Spearman r = 0.6000, P = 0.1750), but are influenced by uncertainty (Kruskal-Wallis H(2) = 10.05, P = 0.0066); n = 10 mice, data are mean +/-SEM. E. Spearman’s correlation of HON-GCaMP max Z-scored fluorescence stimulus values vs VTA DAN-GCaMP values pooled across all reward probabilities. r^2^ = 0.0021158, p = 0.7511. F. Top to bottom: the explained variance of DAN and HON activity (r^2^ score) in our multivariate model for each mouse (Figure 1C), and corresponding contributions of movement, probability and uncertainty. G. Relative contributions of each feature to the model, averaged across the mice.

Fiber photometry does not resolve signals in individual HONs, and may not simply reflect somatic calcium in all cells ^65^. To gain greater resolution of tracking strategies across HONs, we therefore used GRIN lens 2-photon imaging to resolve somatic GCaMP signals of individual HONs ^47^ (Fig. 3A-C); thought to be proportional to their firing rate ^47^. By applying our multivariate encoding model to each HON, we could infer contributions of each predictor to the explained variance of that neuron (Fig. 3D; Fig. S3). Within the investigated features, and in line with photometry recordings, movement was still the dominant contributor to the explained variance in HON activity (Fig. 3E,F). To further quantify whether individual HONs multiplex several features or specialize in encoding movement, we performed unsupervized clustering of the HON cue responses (Fig. 4A). This revealed distributed coding of movement, reward probability, and reward uncertainty across functionally clustered HON subpopulations (Fig. 4B). Some HONs coded for all three features, while others specialized in transmitting information about one feature (Fig. 4C). Overall, however, predominant or considerable movement representation was detected in ∼60% of individual HONs (Fig. 4B,C). Together, these cellular data reveal a dimension of HON information transmission not apparent in photometry analyses, demonstrating functional HON clusters performing specialized coding of different subsets of movement and reward features, but overall predominantly specializing in tracking behavior during the cue period.

**Figure 3.**
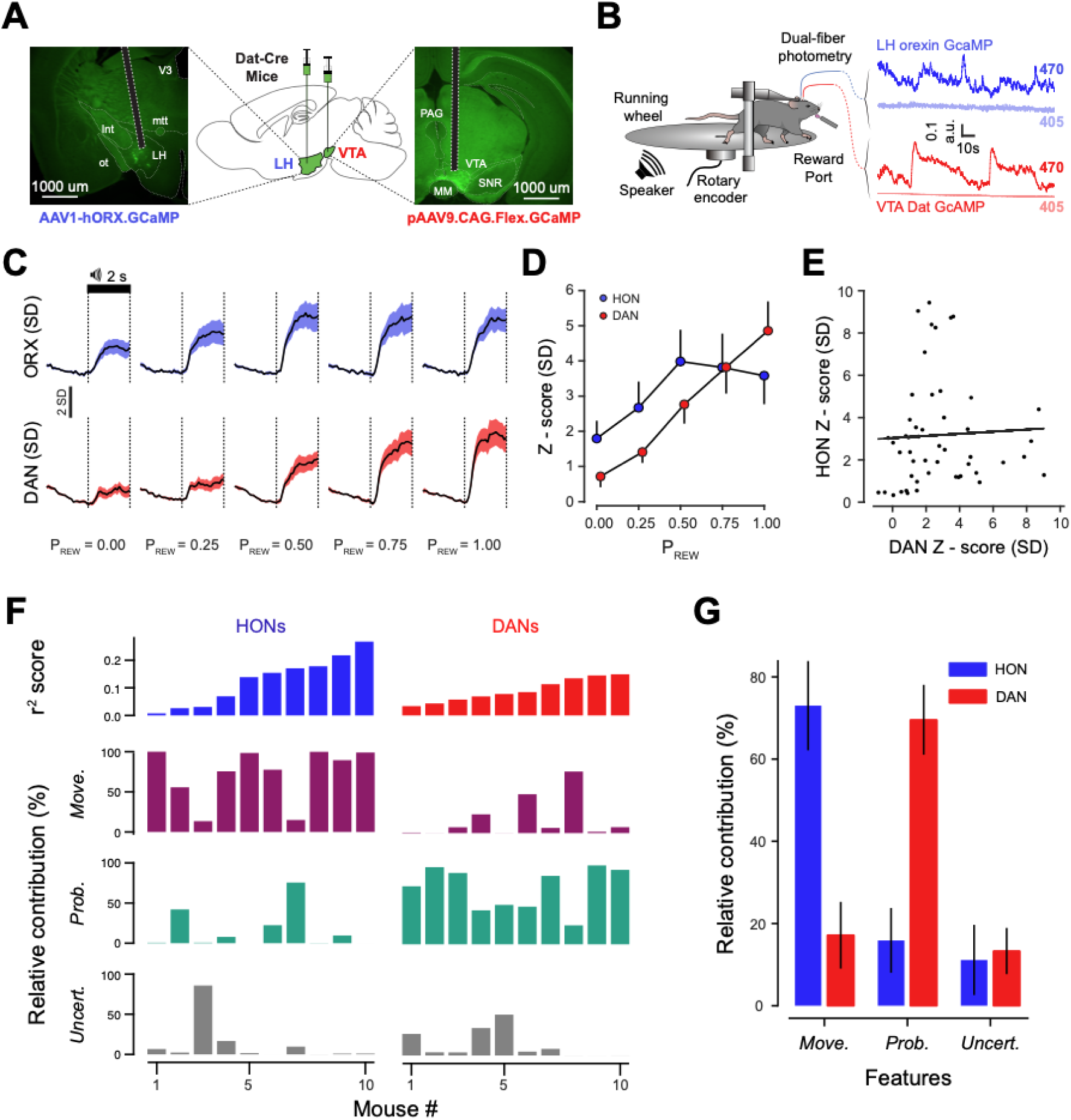
During cue period, some individual HONs multiplex diverse information. A. Left, coronal section example showing HON-GCaMP6s expression and GRIN lens position. Inset, magnified localisation of HON-GCaMP6s expression in LH. Right, behavioral and 2-photon imaging apparatus. B. Left, example two-photon image plane. Right, raw GCaMP fluorescence from example HONs, with accompanying trial and behavioral features. C. Top, heatmaps for HON responses to cues signaling the five reward probabilities. Bottom, Corresponding averages. n = 1197 cells from n = 4 mice. D. Top to bottom, the explained variance of each cell’s activity in the multivariate model (Fig. 1C), and corresponding contributions of movement, probability and uncertainty. E. Relative contributions of each feature to the model, averaged across HONs. F. Distributions of cell contributions for each of the investigated features.

**Figure 4.**
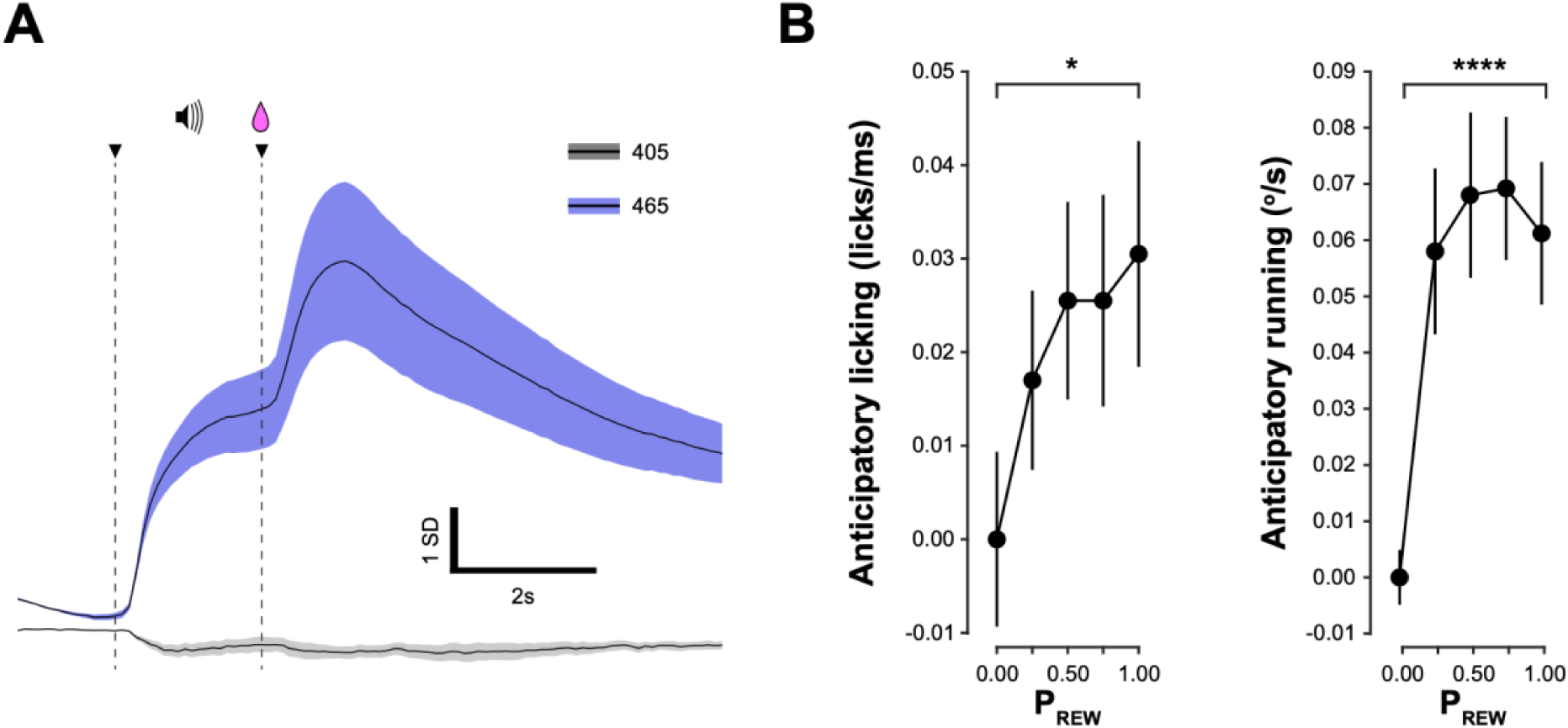
Movement information across the functional clusters of HONs during the cue period.|. A. GMM clustering of relative contribution of HONs to uncertainty, probability and movement features. Bayesian Information Criterion (BIC) score was used to determine the optimal number of HON clusters. B. Proportion of HONs in each HON cluster identified in (A). C. Average relative contribution to uncertainty, probability and movement features of HON clusters identified in (A).

### During the outcome period, HONs primarily track reinforcement identity

Next, we looked into HON activation during the outcome period of the associative cue-reward pairing (Fig. 1A). As before, we first evaluated the concurrent HON and DAN photometry recordings (same set up and experiments as in Fig. 1A and Fig. 2A,B). In simple bivariate analyses, DAN responses to reward displayed an inverse relationship with reward probability, i.e. ‘reward prediction error’^1^, but HON responses to reward were invariant across reward probabilities (Fig. 5A-C). This implies that at the population level, HON signal transmits information not about reward prediction error, but simply about presence or lack of reward, and/or about behavior accompanying reward consumption. Multivariate analysis (Fig. 1B) clarified that reward presence or absence, rather than behavior, was the primary feature explaining HON signal variance in the outcome period (Fig. 5D-E: DANs also displayed such reward coding in addition to reward prediction error, as expected ^54^). The latter conclusion about HONs was confirmed when different behavioral elements (licking, running) were analyzed separately (Fig. S5A).

**Figure 5.**
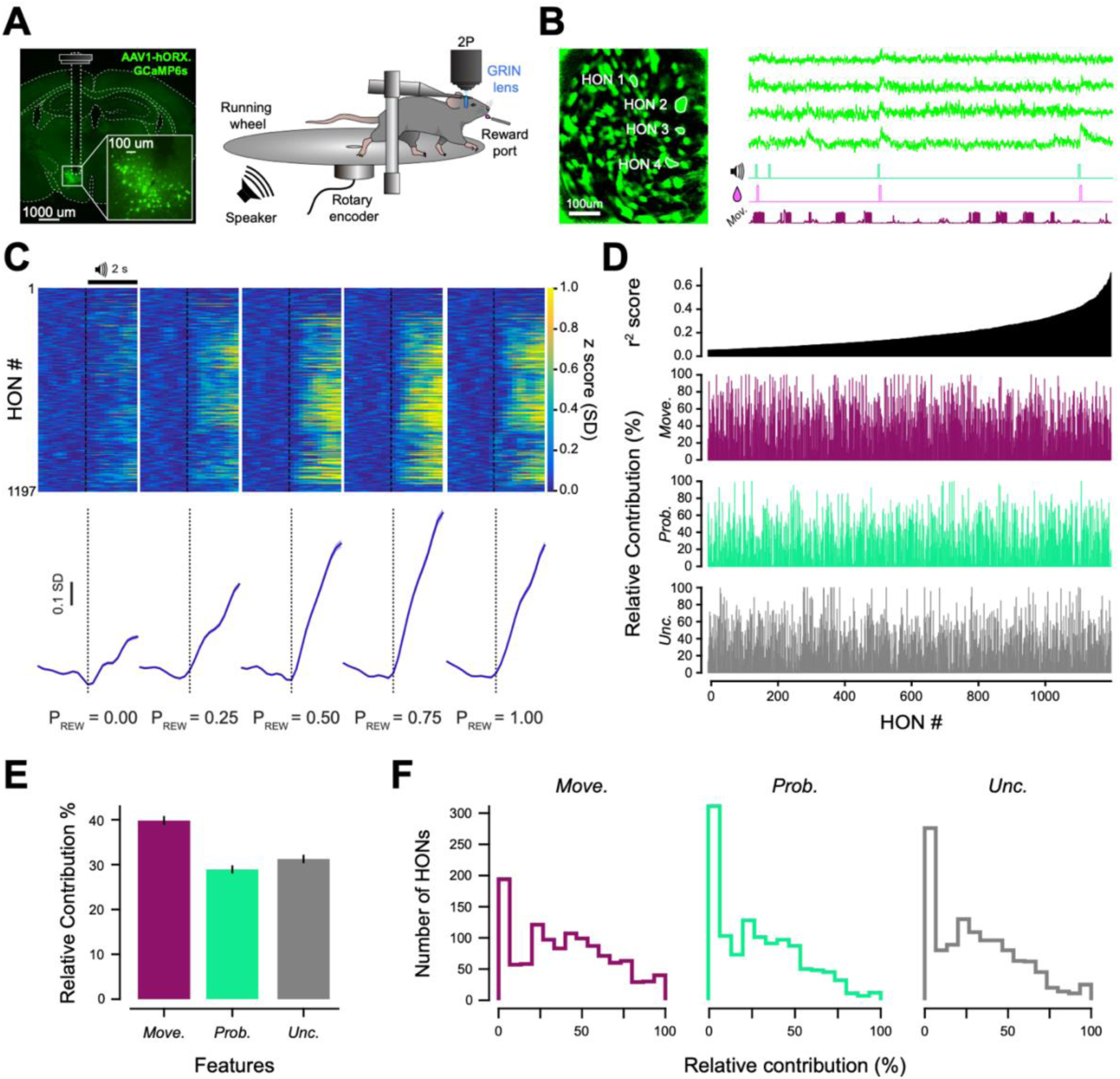
During outcome period, HON population primarily tracks reinforcement identify. A. Responses of LH HON-GCaMP and VTA DAN-GCaMP signals during rewarded (solid lines) and unrewarded (dashed lines) trials, across the five reward probabilities. Each trace is a mean of n = 7 mice. B. Same as (A), but with reward responses visualized more selectively, by subtraction of corresponding unrewarded trial traces. Data are means +/-SEM of n = 7 mice. C. Summary of data in B for maximal responses. DAN responses monotonically decreased with reward probability, i.e. they coded reward prediction error as previously described^1^ (Spearman r = -0.5143, P < 0.001), but HON responses did not monotonically follow reward probability (Spearman r = 0.1266, P = 0.1286), n = 7 mice, data are mean +/-SEM. D. Top to bottom: the explained variance of DAN and HON activity in our multivariate model for each mouse (Figure 1C), and corresponding contributions of movement, probability and uncertainty. E. Relative contributions of each feature to the model, averaged across the mice.

Switching to 2-photon recordings to gain single-cell resolution of the HON responses, we found that HON activation during the outcome period was more homogenous than during the cue period (Fig. 6A,B), with reward being the leading contributor to explaining HON signal variance (Fig. 6C,D; Fig. S4). Similarly to photometry data (Fig. S5A), splitting the behavior feature into licking and running had no effect on the prominence of the reward information in HONs (Fig. SB), confirming that HONs were tracking reward outcome, rather than licking behavior associated with reward consumption. Unsupervised clustering of the HON outcome period responses confirmed this conclusion (Fig. 6E-G). In addition to the reward-specializing HONs (∼52.8% of cells), we observed a smaller fraction of cells whose responses to reward primarily co-varied with reward uncertainty (∼11% of cells) (Fig. 6F,G). We observed no evidence that the functional HON clusters identified during cue or outcome periods were spatially distinct in the LH (Fig. S6).

**Figure 6.**
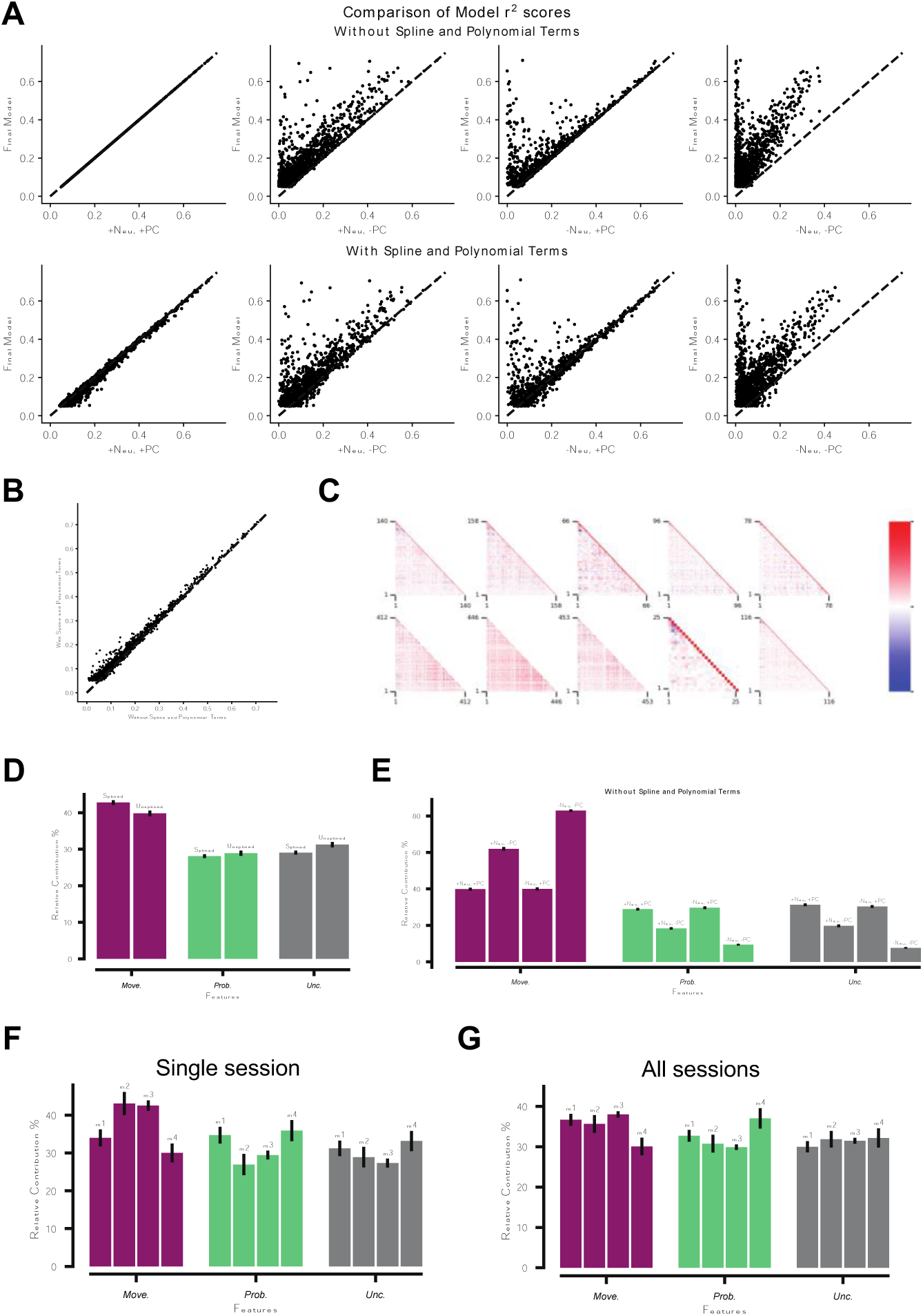
Reinforcement information across the functional clusters of HONs during the outcome period. A. Top to bottom: the explained variance of each HON’s activity in the multivariate model (Fig. 1C), and corresponding contributions of reward, movement, and reward probability and uncertainty, for the reward outcome period. B. Cross-correlations of neural activity across HONs (each point corresponds to a recording session, n = 10 sessions from 4 mice). C. Relative contributions of each feature to the model, averaged across the cells. D. Distributions of cell contributions for each of the investigated features. E. GMM clustering of relative contribution of HONs to reward, movement, uncertainty, probability and movement features. Bayesian Information Criterion (BIC) score was used determine the optimal number of HON clusters. F. Proportion of HONs in each HON cluster identified in (E). G. Average relative contribution to uncertainty, probability, movement and reward features of HON clusters identified in (F). Cluster names and colors as in E.

Overall, these data indicate that during the outcome period, a dominant functional HON cluster performs simple coding of reward vs lack of reward.

## DISCUSSION

Bringing together a multi-dimensional cognitive paradigm, HON neural recordings at both single-cell and population levels, and a multivariate encoding model that enables the contributions of distinct features to be disentangled, we identified specific features associated with HON activity during evolutionarily critical cue-reward associations. In both population (photometry) and individual (single-cell 2P) HON recordings, our data suggest that, during the cue period, HONs primarily specialize in tracking body movement, while in the outcome period, they primarily track the binary presence or absence of reward. This approach highlights the importance of recording body movements and multivariate analyses in cognitive tasks, and suggest a simple functional logic of HON activation during associative perception: HON-transmitted arousal is primarily allocated to tracking actions and reinforcements.

Despite the simple logic of HON activation at the population level during the cue period (Figs. 1G, 5E, S5A), a small subset of HONs deviated from this logic (Fig. 4B,C), and appeared to track multiplexed cognitive information about reward uncertainty or probability. Encoding the latter is a canonical function of VTA DANs ^60, 66, 67^, confirmed here with both simple and multivariate analyses. Without knowing the sequence of activation between HONs and DANs, it is difficult to speculate the function of this redundancy in encoding the same information. One hypothesis may be that, during the cue-outcome period, subpopulations of HONs receive information from DANs ^68^ about the probability or uncertainty of a reward occurring, allowing them to modulate arousal levels appropriately.

During the reward outcome period, more HONs were recruited to signal the presence or absence of reward (Fig. S2A, 5A, Fig. 6B). However, HON amplitude was maintained across reward probabilities (Fig 5B). Furthermore, multivariate analyses suggested that reward-associated body movements, including the ingestive behaviour of licking, were less important than the reward outcome. This suggests HONs transmit a simple binary logic to downstream areas about the nature of reinforcement. As with the cue period, this may allow HONs to allocate appropriate (i.e. increased) arousal during the reward period, potentially enabling downstream systems to form associations more easily. This could potentially serve as a start point from which associative systems can track back in time to identify events preceding reinforcements, helping to improve the efficiency of cue-outcome association formations. Given the importance of HONs in both energy balance^40^ ^43, 69–71^ and addiction ^44, 72–75^, it would be interesting in future to determine if this binary logic is maintained when reward size is altered, if the animal’s energy state were depleted further, or in the presence of substances of abuse.

Our data open multiple additional directions for further studies. Upstream, among the highly diverse direct neural inputs of HONs ^46^, it will be important to determine the sources of motor and reward information during cue and outcome periods, as well as their role in information-gathering of the slower metabolic variables (nutrients and hormones) ^41^, ^76^ ^77^ ^78^ ^79^ ^40^. On a whole-brain level, it remains to be determined how HONs interact other neural systems implicated in associative information processing, such as DANs as well as other subcortical aminergic systems ^80^, to create survival-relevant interpretation of salient events and their behavioral context. Together with axon projection and molecular complexities of HONs ^33, 81–101^, the information specialization revealed here may aid the CNS to interpret the external and internal information they transmit, and increase our understanding of brain disorders involving HONs ^92, 102–108^.

## METHODS

### Animals and gene targeting

All animal procedures were performed in accordance with the Animal Welfare Ordinance (TSchV 455.1) of the Swiss Federal Food Safety and Veterinary Office, and were approved by the Zurich Cantonal Veterinary Office; or approved by the UK Home Office. Mice were kept on a reversed 12-h/12-h light/dark cycle and water, and unless otherwise stated, chow ad libitum. Experiments were performed during the dark phase. All mice were over 8 weeks old. WT C57BL6N mice were used for 2-photon recordings of orexin neurons, and DAT-Cre mice (Jax: Slc6a3tm1(cre)Xz) were used for dual photometry recordings from orexin and dopamine neurons. For recording of LH orexin/hypocretin cell activity, we used AAV1-hORX.GCaMP6s (2.0 x 10^13^ vg/ml, Penn vector core). This virus targets orexin/hypocretin neurons with >96% specificity ^49^. For recording VTA dopamine cell activity, we used pAAV9.CAG.Flex.GCaMP6s.WPRE.SV40 (1.9 x 10^13^ vg/ml, Addgene) or pGP-AAV1-syn-FLEX-jGCaMP7f-WPRE (titre 2.1 x 10^13^ vg/ml, Addgene). 6s and 7f dopamine recordings were pooled in presented analysis, because their response amplitudes were statistically indistinguishable.

### Surgeries

Mice were anesthetized with isoflurane, placed in a stereotaxic frame (Kopf instruments), and a small incision was made to expose the skull surface.

For fiber photometry implants in the lateral hypothalamus and the ventral tegmental area, two small craniotomies were drilled at AP: -1.35, ML: +/-0.9 (to target the LH) and AP: -3.32 and ML: +/-0.5 (to target the VTA). A Nanoject III (Drummond Scientific) was used to deliver three 50nL injections of AAV1-hORX.GCaMP6s in to the LH at a rate of 1nL/s, with coordinates from bregma of AP: -1.35, ML: +/-0.9, DV: -5.7, 5.4 and -5.1, and two doses of 50nL flex-GCaMP6s or 7f were injected into the VTA in the contralateral hemisphere, with coordinates from bregma of AP: -3.32, ML: +/-0.5, DV: - 4.4 and -4.2. The bone surface was roughened using a 0.8 mm diameter dental drill to remove periosteum and promote adhesion, and a custom stainless steel headplate (Protolabs) was glued to the skull surface using clear Histoacryl (Braun). Optic cannulas with a 200 μm core fiber optic (ThorLabs CFML12U-20, NA 0.39) in custom-built holders were then advanced to a depth 50 - 100 μm above the most superficial viral injection site in each brain region. One of the two fibers was angled at 10 degrees to make space for securing optical cables later. This was randomized between VTA and LH. The optic-fiber cannulas and headplate were then secured using three-part dental cement (Superbond, Sun Medical), and the fiber holders retracted. The cement was coated in black nail varnish (Essie), and the skin glued using Histoacryl around the implant site.

For implanting GRIN lenses in the lateral hypothalamus, a 1.2 mm craniotomy was drilled at AP: -1.35 mm, ML: +/-0.9 mm. A Nanoject III (Drummond Instruments) with a glass micropipette was used to inject three 50 nL injections of hORX.GCaMP6s at a rate of 1nL/s, with coordinates from bregma of AP: -1.35, ML: +/-0.9, DV: -5.7, 5.4 and -5.1. After removal of the injection pipette, the GRIN lens (7.3mm long, 0.6mm diameter cylindrical lens, Inscopix), was lowered in the center of the craniotomy to a depth of -5.3 mm from the height at the nearby skull surface using a linear manipulator (Scientifica), at a rate of 2 um/s. The lens was secured using three-part dental cement (Superbond, Sun Medical), a custom head plate was attached to the skull and GRIN lens cap using histoacryl, then three-part cement. The cement was then coated in black nail varnish, and the skin glued using Histoacryl around the implant site.

### Fiber photometry

Mice were head-fixed and allowed to run on a horizontal wheel, attached to a rotary encoder (Figure 1A, 2A), and cells in the LH and VTA of fiberoptic-implanted mice were imaged using fiber photometry similar to our previous work ^109^. Briefly, GCaMP was excited with blue and violet LEDs (465 and 405 nm respectively, Thorlabs), alternating at 10 Hz, and resulting fluorescence emission collected using a photoreceiver (Thorlabs) through a fiber-connectorized GFP filter cube (Doric, FMC_GFP_FC). Average excitation light power was ∼100 μW at the fiber tip. Fluorescence produced by the isosbestic 405-nm excitation was used to check for motion artifacts ^110^ (Figure S2A). Auditory tones and reward delivery were controlled using custom-built Labview software, and outputs from a lick sensor attached to the reward spout, the rotary encoder and photometry signal were acquired at a rate of 400 Hz using a HEKA LH1+1 I/O box, and HEKA Patchmaster software. 465-nm excited signal was calculated by taking the mean of 6 samples centred on the mid-point of the period when 465-light was triggered, and similarly for 405-nm. To produce Z-score values for photometry traces (Figures 2C-E, 5A-C), raw fluorescence values were cut into peri-event windows either side of the stimulus onset, and the mean and standard deviation (SD) of the 2s pre-stimulus baseline was calculated. Each sample was then Z-scored using the following calculation [100*((sample - mean baseline)/SD baseline)] as in (Duffet et al., 2022). These traces were then filtered using a Savitsky-Golay filter and the median of trials for each mouse were used to generate traces for each probability. For the encoding model described below, Z-scored traces were scaled from 0-1.

### 2-photon GRIN lens imaging

Hypothalamic 2-photon imaging was performed similarly to our previous work ^47^. The setup was as described above in the photometry experiments, except the mouse’s head was positioned under the microscope objective (Figure 3A). Changes in GCaMP fluorescence were imaged with a custom-built electro-tunable lens equipped resonant/galvanometer scan head two-photon microscope (INSS) and a femtosecond-pulsed mode-locked Ti:sapphire laser (Spectra-physics Mai Tai HP Deepsee 2) at 950 nm through a 20 x (0.45 NA, Olympus) air-IR objective at 31 frames/s with 512 x 512 pixel resolution, using custom Labview software ^47^. Six planes were imaged using the electro-tunable lens, yielding a volume rate of 5.1 volumes/s. Images were obtained with a 510/80 nm band-pass emission filter. A National Instruments USB-6008 DAQ board was used to output TTL triggers and count imaging frames. Auditory tones and reward delivery were controlled using custom-built Labview software, and outputs from a lick sensor attached to the reward spout and rotary encoder were recorded in the 6008 DAQ board. Frame outputs from the DAQ board were used to trigger trial onsets in Labview software on a separate PC, to ensure stimuli delivered by the USB 6343 (NI) were always aligned to the same plane. Bottom planes were aligned as best as possible to the same depth each session.

Suite2p ^111^ was used for image processing, including correcting motion artifacts and drift in the imaging plane, and detection of regions of interest (ROIs) around HONs. Mean intensity within each ROI was used to generate F(raw). F(raw) traces were cleaned of artifact spikes by excluding values greater or lesser than the mean +/-50 times the standard deviation of neighboring samples. A rolling window of 250 seconds in the photometry data and 20 seconds in the 2-photon data was used for removal of these larger spikes. Traces were excluded if the neuropil activity (F(neu)) and HON activity F(raw) had similar means, i.e the mean difference between F(raw) and F(neu) was less than 100 - a threshold decided by observing the data. Traces were further excluded if they showed no deviations from the baseline above the mean +/-2*sigma boundary in more than 5% of the whole trace. ROIs were grouped into clusters using agglomerative clustering, measuring the spatial distance between the medians of the ROIs. ROIs that had a distance less than 20 pixels apart were grouped together and only the best ROI (with greatest difference between F(raw) and F(neu) was kept. 1990 ROIs, from 10 sessions from 4 animals were detected. The selected F(raw) and F(neu) traces were Z-scored locally as in photometry data above. The traces were then smoothed using a low pass 8^th^ order Butterworth filter before being used in bi- and multivariate analysis.

### Pavlovian conditioning

At least 14 days after surgery, mice were habituated to headfixation and running on the wheel for 2 sessions of around 20 minutes each. In the next 2 sessions of headfixation, they were habituated to auditory stimuli with 15 – 30 trials of each frequency, presented in a random order. Cues were pure tones of 4, 6, 8, 10 and 12 Hz, 2s in duration, at an acoustic power of 60-70 dB, played using the PC soundcard, through a speaker (Visaton Titankalotte DSM 25 FFL, 8 Ohm). Next, mice were food restricted overnight and habituated to reward by positioning the lick spout 2-3 mm from the tongue, and presenting milkshake rewards (Strawberry Energy Milk Drink, Emmi) of 14 ul at random intervals from 5 - 9 seconds, until the mouse stopped licking. Typically this took 10 - 40 trials. Food restriction was maintained from then on, with full access given at weekends. In the next session, mice were presented with the 5 pure tones of 2 s duration in a random order, each tone paired with a given probability of reward, which was delivered at the offset of the auditory tone ^60^. Reward probabilities (PREW) of 0, 0.25, 0.5, 0.75 and 1.0 were paired with tones of 4, 6, 8, 10, 12, or 12, 10, 8, 6, 4 Hz, respectively. Tones were only delivered after no running was detected for 2 s. In a given session, trials were presented until mice stopped licking, typically 70-150 trials per session including 15 - 30 trials per stimulus. In total, mice performed 150 - 200 trials per reward probability over the course of 7 - 8 days. After conditioning, anticipatory movement evoked by the cue increased with the reward probability indicating that mice learnt to behaviourally discriminate the stimuli (Figure S2B). To investigate reward prediction error (RPE, Figure 5A-C), 7 mice were further trained on 1000 trials per cue and a reward was delivered during the outcome period of one P_REW_ = 0 trial once per session every two or three sessions. This resulted in an unexpected P_REW_ = 0 reward rate of around 0.02.

### Bivariate analysis of neural coding

Coding of reward uncertainty was evaluated by quantifying the influence of reward uncertainty on neural activity. To achieve this, we followed the method of ^60^; in each dataset in which the five reward probabilities were tested, the increases in neural activity during the cue (Figures 1 - 3) or outcome period (Figure 4) were ranked across the five probabilities for each cell (in 2-photon experiments) or each mouse (in photometry experiments). A Kruskal-Wallis test was then performed on the ranked values, with three groups defined by the degree of uncertainty (i.e. p=0.0 and 1.0; p=0.25 and 0.75; p=0.5). The ranking of the data points accounted for the paired nature of the data from each mouse or neuron, while the Kruskal-Wallis test is appropriate for multiple comparisons of nonparametric data. Coding of reward probability was evaluated by quantifying monotonic relationships between reward probability on neural activity ^60^. The strength and direction of monotonic relationships between probability and neural activity was assessed with Spearman’s correlation analysis. The statistical analyses were performed using Prism 8.4.3 (GraphPad). In Figure 2D, E, asterisks represent P values from Bonferroni post-hoc analysis following RM ANOVA (ns = P>0.05, *P<0.05, **P<0.01, ****P<0.0001); main ANOVA results are reported in corresponding Results text.

### Multivariate Analysis

An encoding model based on a previous study ^54^ was then applied, and involved multiple regression analysis with neural signals (photometry or 2P GCaMP recordings) as the target feature, and movement, reward, reward probability, and reward uncertainty as input features (Figure 1C). The movement feature was a linear average of lickometer and running sensor data. Additional regressions were performed with these features as separate inputs, which produced similar conclusions (data available on request). Features were downsampled to match the respective sampling rates of photometry and 2-photon neural activity recordings. The probability feature contained the 5 levels of probability (Figure 1A). The uncertainty feature contained 3 levels and was computed from probability based on ^60^ (low uncertainty, P_REW_ = 0 and 1; medium uncertainty, P_REW_ = 0.25 and 0.75, high uncertainty, P_REW_ = 0.5). The reward feature was coded as 0 or 1 for the full duration of the outcome window depending on the presence or absence of reward for each trial. The cue and outcome periods were modeled separately. For each window, the features and neural activity were similarly segmented and scaled from 0 to 1 to keep the range consistent across features and targets.

For two-photon data, the neuropil activity of each HON trace was processed similarly to the target neural activity and fed into the model to improve its performance. For each trace in a mouse, principal component analysis (PCA) was performed on all other traces pertaining to that animal and the first principal component (PC) was also given as an input into the model, based on ^54^. To incorporate non-linear terms into the model, the behavioral features were squared and used along with the input features. The behavioral and trial features (except Reward) were convolved with a spline basis of polynomial degree 3, using the sklearn SplineTransformer in Python, to add continuity and allow for lag caused by GCaMP. The model was regularized with a ridge regression model to account for the multi-collinearity of the input features. HON traces where the r^2^ score is less than 5% were excluded from further analyses and the figures. 1197 and 1666 traces were ultimately used for analyses of the cue and outcome period respectively, as indicated in Figures 3, 4 and 6. The models were fitted with a 5-fold cross-validation and the average r^2^ score of the model was calculated, ignoring any score below 0. The final model used for analyses (in Figure 2 and 4) included splined and polynomial terms along with the neuropil and 1^st^ PC traces. A comparison of the r^2^ scores of the final model against models without each of the features listed above can be found in Figure S3 (A-B) for the cue period and S4(A-B) for the outcome period. No major differences were observed in the percent of relative contributions across cells when taken across the different models, as can be observed in Fig S3 (D-G) for the cue period and Fig S4(D-G) for the outcome period. The analysis pipeline is shown in Figure S1.

The contribution of each feature was quantified by observing how the performance of the model changed when each feature was excluded (based on ^54^). For a combination of features, we first calculated the r^2^ score of the full model with all the features. From this, we recursively excluded one feature at a time and calculated the new r^2^ score of the partial model. The relative contribution of the feature to the model is the relative decrease in the r^2^ score of the partial model compared to the full model ^54^. The relative contributions were calculated for each trace and averaged across the traces for each feature to investigate the importance of the features to the HON population, as in Figures 3E, S3(D-G), 6C and S4(D-G).

### Cluster Analysis

A Gaussian Mixture Model (GMM) was used to perform unsupervized clustering of the relative contributions of features to the model for each HON trace, which identified different functional sub-populations of HONs. The number of clusters chosen for clustering was decided by the Bayesian Information Criterion (BIC) and the ‘elbow method’ was employed to choose the optimal number of clusters, wherein increasing the number of clusters did not significantly improve the BIC score as show in Figures 4A and 6E. Spherical clusters were used due to their robustness to outliers and interpretability.

### Histology and Immunohistochemistry

Mice were terminally anaesthetized and perfused transcardially with 4% PFA. Brains were post-fixed for 24 h, placed in 30% sucrose for a further 24 hours, then frozen in dry ice and stored at -80°c. 50 um coronal brain slices were later sectioned using a cryostat. Epifluorescence images were then acquired and stitched using an epifluorescence microscope (Nikon) and merged in ImageJ.

## ACKNOWLEDGEMENTS

This work was supported by ETH Zürich. We would like to thank Mathilde Guillaumin and Alexander Tesmer for help with statistical analysis, Paulius Viskaitis for help with initial setting up of experiments, and Cristina Concetti for help with some of the surgeries.

## AUTHOR CONTRIBUTIONS

DB conceived the study, EB and DB conceptualized and designed the experiments. EB performed the experiments, and analyzed the data. AA designed and validated the multivariate encoding model, and performed the multivariate analyses. NG contributed to experiments and analysis. All authors contributed to data interpretation and the writing of the manuscript.

---

**Supplementary Figure S1.**
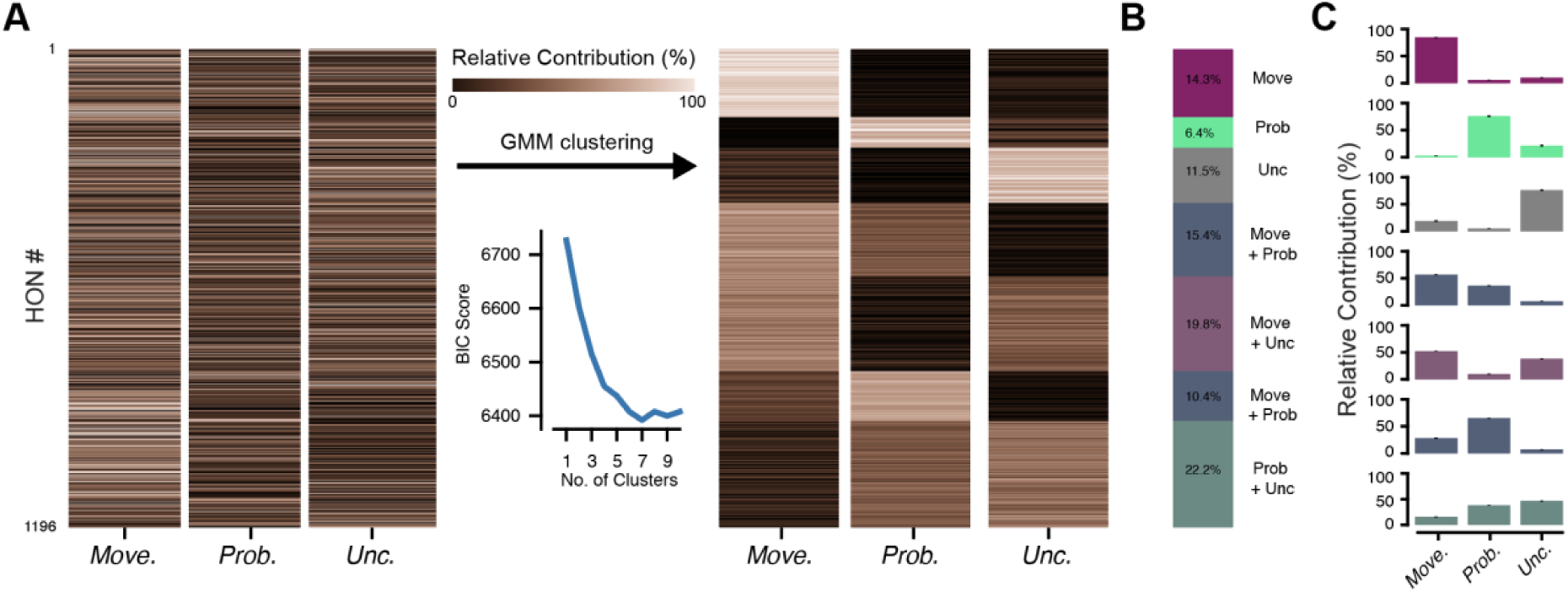
Data analysis pipeline for multivariate analysis. Flow-chart of data analysis pipeline (the example shown is for the 2-photon imaging experiments), as further described in the Methods section.

**Supplementary Figure S2.**
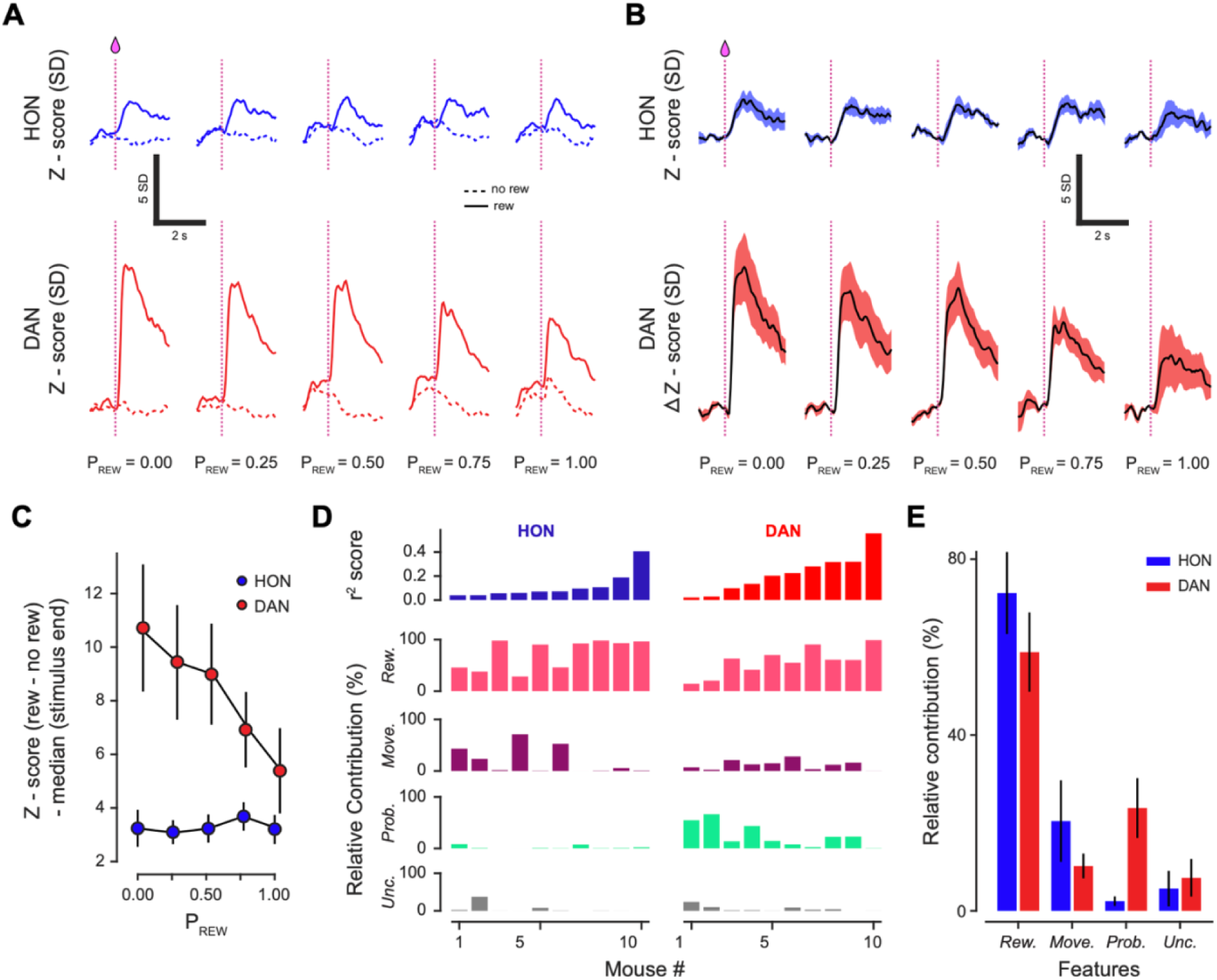
Control experiments: movement artefacts and behavioral confirmation of Pavlovian conditioning. A. Mean Z-scored photometric fluorescence HON-GCaMP values during cue and reward outcome presentation, averaged across all trials and all reward probabilities, n = 10 mice. Fluorescence recorded at 465nm (blue) and 405nm (black) shows calcium-dependent, and movement-related changes respectively. B. Example anticipatory licking (left), and running (right) averaged across 50 trials and five probabilities during stimulus presentation. Rank sum of P_REW_ 0 vs P_REW_ 1 is shown, *, p < 0.02, ****, p < 0.0001. All mice had to demonstrate a significant difference between P_REW_ 0 and P_REW_ 1 to be included in the study.

**Supplementary Figure S3.**
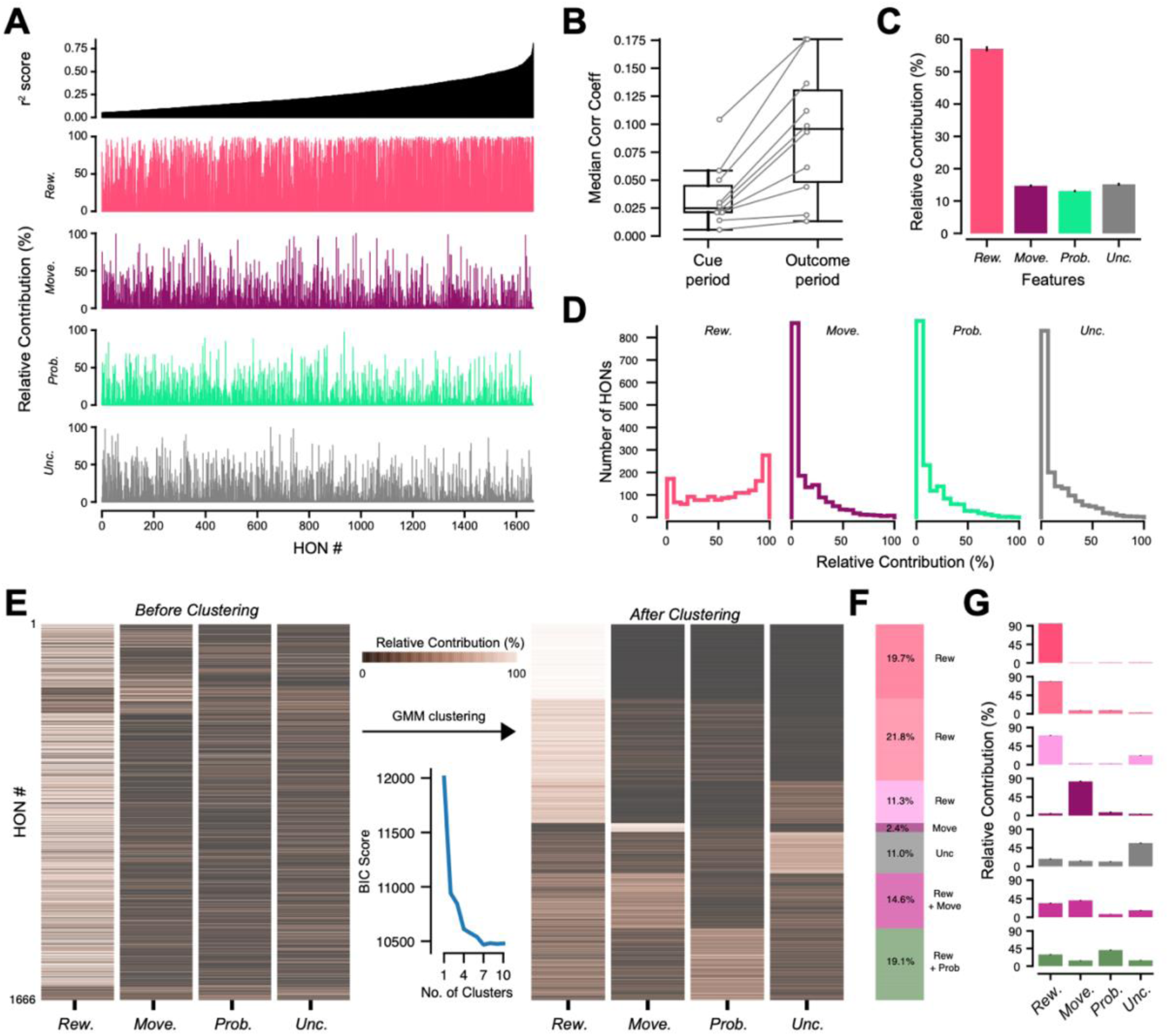
Model evaluations and comparisons (Cue period, 2-photon data) A. Comparison of r^2^ scores of models with different combinations of non-feature inputs in the cue period. ‘Final model’ refers to the model used to present results in the paper, with input features remaining un-splined and no polynomial features added. The final model includes the neuropil (*Neu*) and first principal component (*PC*) and the plots show the differences when these two non-input features are included or excluded (*+* or *-*). Top, models have no polynomial features or splined input features. Bottom, with behavioral features splined and polynomial features added. See Methods for details. B. Comparison of r^2^ scores of models with and without splined input features and polynomial terms highlighted from A. C. Heatmap of correlation coefficient values between cell traces of concatenated cue windows across all (n=10) sessions per mouse (n=4). D. Comparison of relative contributions of models with and without splined input features and polynomial terms as in B. E. Comparison of relative contributions of models with different combinations of non-feature inputs as in A (Top row). F. Comparison of relative contributions of models from one session from each mouse with the highest number of detected cells. G. Comparison of relative contributions of models from all sessions from each mouse.

**Supplementary Figure S4.**
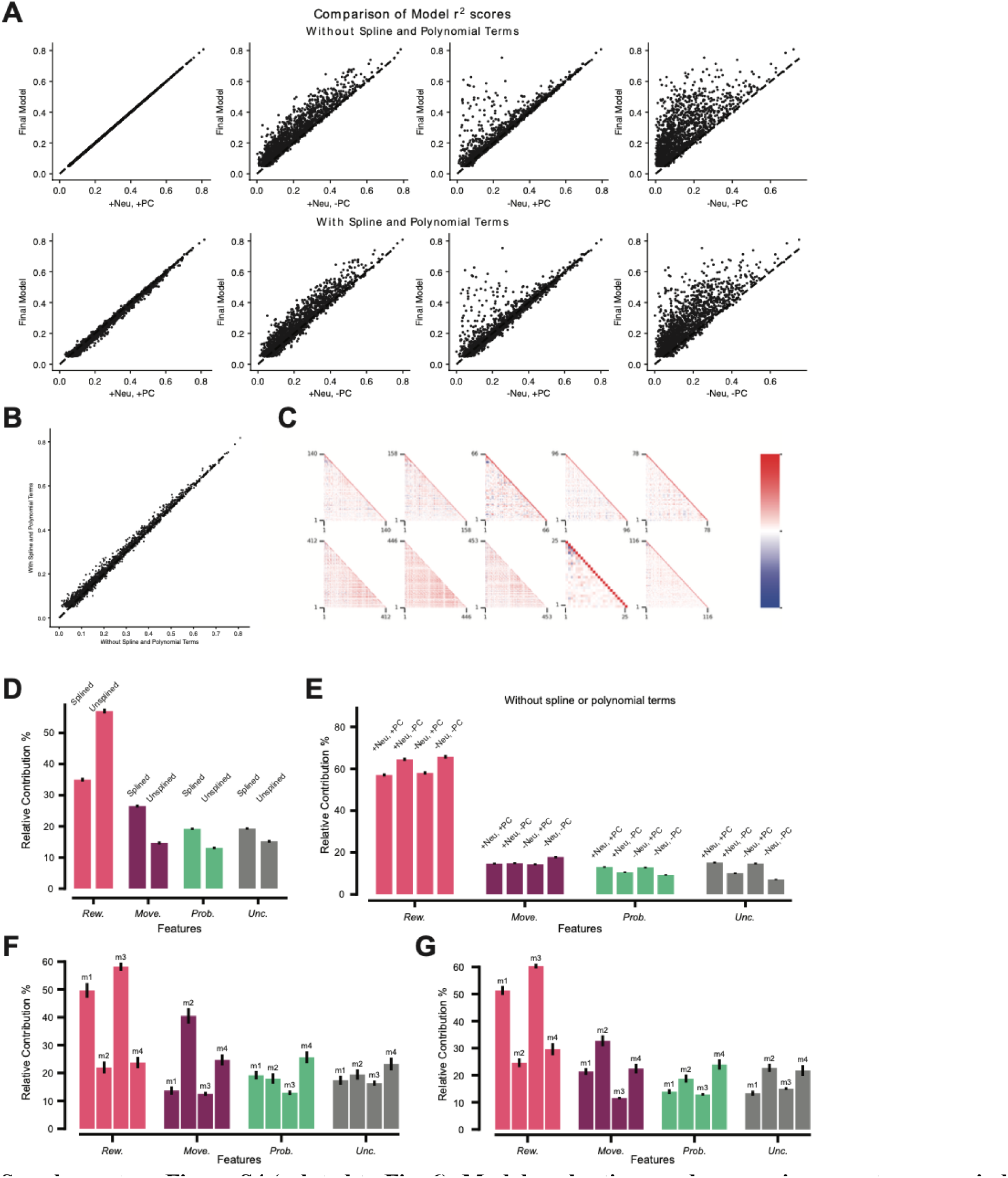
(related to Fig. 6). Model evaluations and comparisons; outcome period, 2-photon data. A. Comparison of r^2^ scores of models with different combinations of non-feature inputs in the outcome period. ‘Final Model’ refers to the model used to present results in the paper, with input features remaining un-splined and no polynomial features added. The Final Model includes the neuropil (*Neu*) and first principal component (*PC*) and the plots show the differences when these two non-input features are included or excluded (*+* or *-*). Top, Models have no polynomial features or splined input features. Bottom, Behavioral features are splined and polynomial features are added. See Methods for details. B. Comparison of r^2^ scores of models with and without splined input features and polynomial terms highlighted from A. C. Heatmap of correlation coefficient values between cell traces of concatenated outcome windows across all (n=10) sessions per mouse (n=4). *Comparison of relative contributions across models:* D. Comparison of relative contributions (%) of models with and without splined input features and polynomial terms. E. Comparison of relative contributions (%) of models with different combinations of non-feature inputs. F. Comparison of relative contributions (%) of models from one session from each mouse with the highest number of detected cells G. Comparison of relative contributions (%) of models from all sessions of each mouse.

**Supplementary Figure S5.**
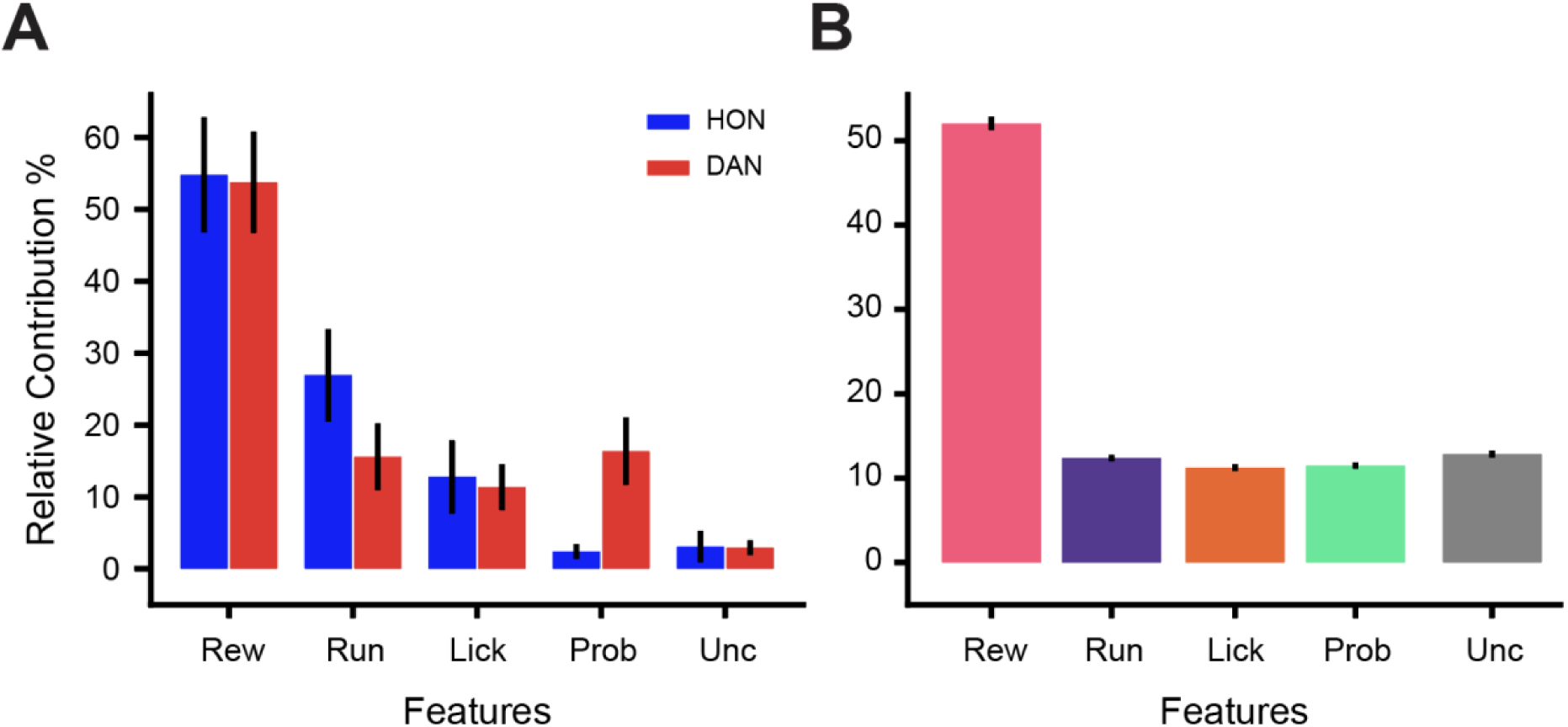
Tracking of HONs to licking and running features during reward outcome period. A. Relative contributions of each feature to the model for photometry during the reward outcome period, averaged across the mice. Data as in Fig. 5E, with movement feature split into licking and running. B. As in A for 2-photon data.

**Supplementary Figure S6.**
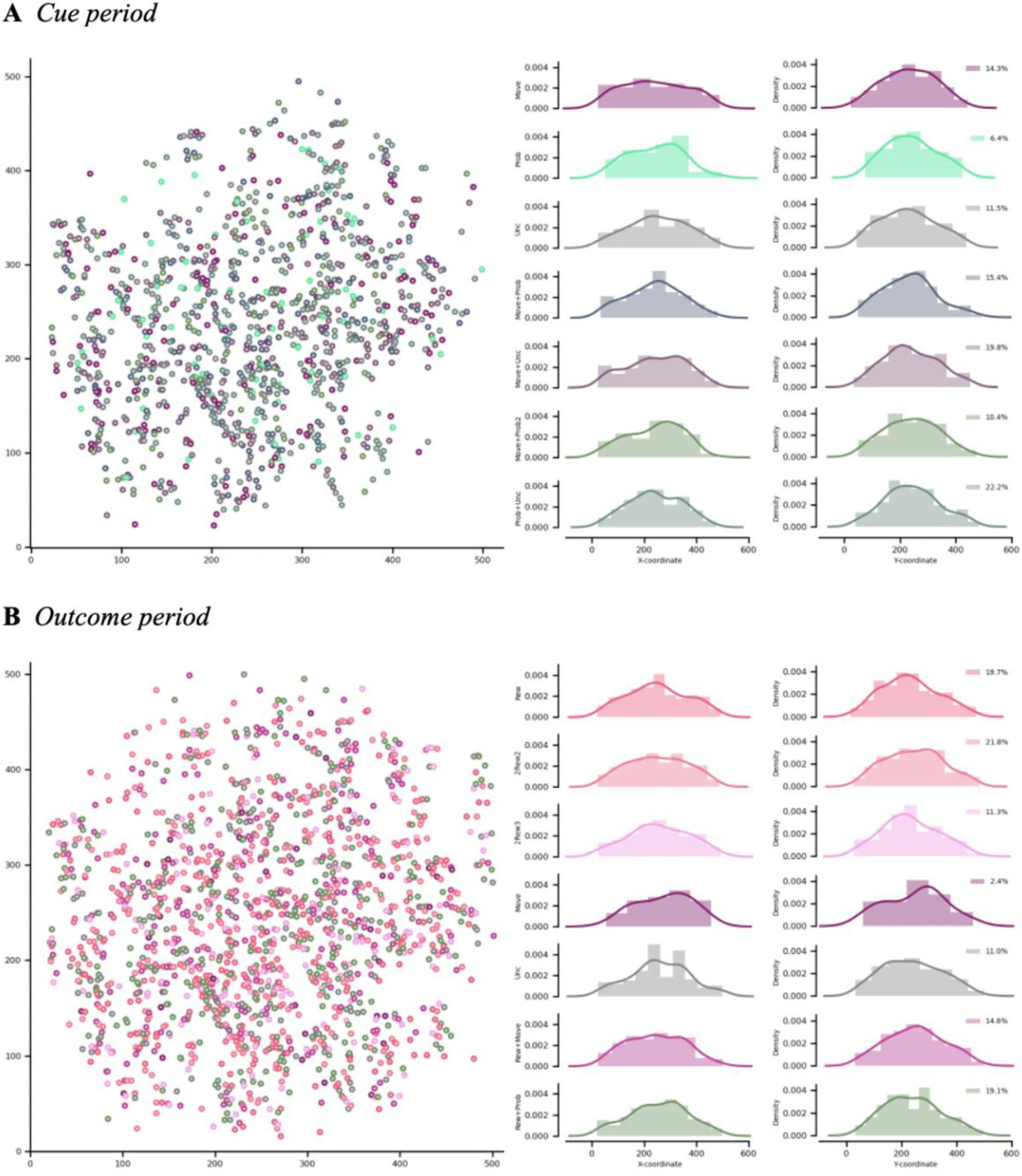
(related to functional clustering in Figs. 4 and 6) Spatial organization of functional HON clusters during cue and outcome periods. A. Locations of identified cells within the frame of the two-photon images (n=512 x 512) color coded according to the clusters identified in Fig 4. B. Density plots of the locations of the cells across the x-axis for each identified cluster C. Density plots of the locations of the cells across the y-axis for each identified cluster D. Locations of identified cells within the frame of the two-photon images (n=512 x 512) color-coded according to the clusters identified in Fig 6. E. Density plots of the locations of the cells across the x-axis for each identified cluster. F. Density plots of the locations of the cells across the y-axis for each identified cluster.

## REFERENCES

1. Schultz, W., Dayan, P. & Montague, P.R. A neural substrate of prediction and reward. Science 275, 1593–1599 (1997).

2. Nassar, M.R., et al. Rational regulation of learning dynamics by pupil-linked arousal systems. Nature neuroscience 15, 1040–1046 (2012).

3. Browning, M., Behrens, T.E., Jocham, G., O’Reilly, J.X. & Bishop, S.J. Anxious individuals have difficulty learning the causal statistics of aversive environments. Nature neuroscience 18, 590–596 (2015).

4. Bar, M. The proactive brain: using analogies and associations to generate predictions. Trends Cogn Sci 11, 280–289 (2007).

5. Clark, A. Whatever next? Predictive brains, situated agents, and the future of cognitive science. Behav Brain Sci 36, 181–204 (2013).

6. Schultz, W. Multiple reward signals in the brain. in Nature Reviews Neuroscience 199–207 (2000).

7. O’Doherty, J.P. Reward representations and reward-related learning in the human brain: insights from neuroimaging. Current opinion in neurobiology 14, 769–776 (2004).

8. Dreher, J.C., Kohn, P. & Berman, K.F. Neural coding of distinct statistical properties of reward information in humans. Cereb Cortex 16, 561–573 (2006).

9. Keller, G.B. & Mrsic-Flogel, T.D. Predictive Processing: A Canonical Cortical Computation. Neuron 100, 424–435 (2018).

10. Livneh, Y. & Andermann, M.L. Cellular activity in insular cortex across seconds to hours: Sensations and predictions of bodily states. Neuron 109, 3576–3593 (2021).

11. Zimmerman, C.A., et al. Thirst neurons anticipate the homeostatic consequences of eating and drinking. Nature 537, 680–684 (2016).

12. Wolpert, D.M. & Ghahramani, Z. Computational principles of movement neuroscience. Nature neuroscience 3 Suppl, 1212–1217 (2000).

13. Rao, R.P. & Ballard, D.H. Predictive coding in the visual cortex: a functional interpretation of some extra-classical receptive-field effects. Nature neuroscience 2, 79–87 (1999).

14. Engel, A.K., Fries, P. & Singer, W. Dynamic predictions: oscillations and synchrony in top-down processing. Nature reviews. Neuroscience 2, 704–716 (2001).

15. Volkow, N.D., et al. Addiction: decreased reward sensitivity and increased expectation sensitivity conspire to overwhelm the brain’s control circuit. Bioessays 32, 748–755 (2010).

16. Dayan, P. & Niv, Y. Reinforcement learning: the good, the bad and the ugly. Current opinion in neurobiology 18, 185–196 (2008).

17. Hassabis, D., Kumaran, D., Summerfield, C. & Botvinick, M. Neuroscience-Inspired Artificial Intelligence. Neuron 95, 245–258 (2017).

18. Munn, B.R., Muller, E.J., Wainstein, G. & Shine, J.M. The ascending arousal system shapes neural dynamics to mediate awareness of cognitive states. Nature communications 12, 6016 (2021).

19. Aston-Jones, G. & Cohen, J.D. An integrative theory of locus coeruleus-norepinephrine function: Adaptive gain and optimal performance. Annual Review of Neuroscience 28, 403–450 (2005).

20. Mileykovskiy, B.Y., Kiyashchenko, L.I. & Siegel, J.M. Behavioral correlates of activity in identified hypocretin/orexin neurons. Neuron 46, 787–798 (2005).

21. Lee, M.G., Hassani, O.K. & Jones, B.E. Discharge of identified orexin/hypocretin neurons across the sleep-waking cycle. Journal of Neuroscience 25, 6716–6720 (2005).

22. Nishino, S., Ripley, B., Overeem, S., Lammers, G.J. & Mignot, E. Hypocretin (orexin) deficiency in human narcolepsy. Lancet 355, 39–40 (2000).

23. Lin, L., et al. The sleep disorder canine narcolepsy is caused by a mutation in the hypocretin (orexin) receptor 2 gene. Cell 98, 365–376 (1999).

24. Peyron, C., et al. A mutation in a case of early onset narcolepsy and a generalized absence of hypocretin peptides in human narcoleptic brains. Nature medicine 6, 991–997 (2000).

25. Chemelli, R.M., et al. Narcolepsy in orexin knockout mice: molecular genetics of sleep regulation. Cell 98, 437–451 (1999).

26. Hara, J., et al. Genetic ablation of orexin neurons in mice results in narcolepsy, hypophagia, and obesity. Neuron 30, 345–354 (2001).

27. Sakurai, T., et al. Orexins and orexin receptors: A family of hypothalamic neuropeptides and G protein-coupled receptors that regulate feeding behavior. Cell 92, 573–585 (1998).

28. de Lecea, L., et al. The hypocretins: hypothalamus-specific peptides with neuroexcitatory activity. Proceedings of the National Academy of Sciences of the United States of America 95, 322–327 (1998).

29. Adamantidis, A.R., Zhang, F., Aravanis, A.M., Deisseroth, K. & de Lecea, L. Neural substrates of awakening probed with optogenetic control of hypocretin neurons. Nature 450, 420–424 (2007).

30. Grujic, N., Tesmer, A., Bracey, E., Peleg-Raibstein, D. & Burdakov, D. Control and coding of pupil size by hypothalamic orexin neurons. bioRxiv https://doi.org/10.1101/2022.04.12.488026 (2022).

31. Harris, G.C. & Aston-Jones, G. Arousal and reward: a dichotomy in orexin function. Trends in neurosciences 29, 571–577 (2006).

32. Hagan, J.J., et al. Orexin A activates locus coeruleus cell firing and increases arousal in the rat. Proceedings of the National Academy of Sciences of the United States of America 96, 10911–10916 (1999).

33. Horvath, T.L., et al. Hypocretin (orexin) activation and synaptic innervation of the locus coeruleus noradrenergic system. The Journal of comparative neurology 415, 145–159 (1999).

34. Sutcliffe, J.G. & de Lecea, L. The hypocretins: setting the arousal threshold. Nature reviews. Neuroscience 3, 339–349 (2002).

35. Kayaba, Y., et al. Attenuated defense response and low basal blood pressure in orexin knockout mice. Am J Physiol Regul Integr Comp Physiol 285, R581–593 (2003).

36. Bonnavion, P., Jackson, A.C., Carter, M.E. & de Lecea, L. Antagonistic interplay between hypocretin and leptin in the lateral hypothalamus regulates stress responses. Nature communications 6, 6266 (2015).

37. Yamashita, A., et al. Aversive emotion rapidly activates orexin neurons and increases heart rate in freely moving mice. Mol Brain 14, 104 (2021).

38. Burdakov, D. Reactive and predictive homeostasis: Roles of orexin/hypocretin neurons. Neuropharmacology 154, 61–67 (2019).

39. Schöne, C. & Burdakov, D. Orexin/Hypocretin and Organizing Principles for a Diversity of Wake-Promoting Neurons in the Brain. Curr Top Behav Neurosci 33, 51–74 (2017).

40. Yamanaka, A., et al. Hypothalamic orexin neurons regulate arousal according to energy balance in mice. Neuron 38, 701–713 (2003).

41. Burdakov, D. Physiological Changes in Glucose Differentially Modulate the Excitability of Hypothalamic Melanin-Concentrating Hormone and Orexin Neurons In Situ. in Journal of Neuroscience 2429–2433 (2005).

42. Cai, X.J., et al. Hypothalamic orexin expression: modulation by blood glucose and feeding. Diabetes 48, 2132–2137 (1999).

43. Viskaitis, P., et al. Ingested non-essential amino acids recruit brain orexin cells to suppress eating in mice. Current biology : CB 32, 1812–1821 e1814 (2022).

44. Boutrel, B., et al. Role for hypocretin in mediating stress-induced reinstatement of cocaine-seeking behavior. Proceedings of the National Academy of Sciences of the United States of America 102, 19168–19173 (2005).

45. Giardino, W.J., et al. Parallel circuits from the bed nuclei of stria terminalis to the lateral hypothalamus drive opposing emotional states. Nature neuroscience 21, 1084–1095 (2018).

46. Gonzalez, J.A., Iordanidou, P., Strom, M., Adamantidis, A. & Burdakov, D. Awake dynamics and brain-wide direct inputs of hypothalamic MCH and orexin networks. Nature communications 7, 11395 (2016).

47. Karnani, M.M., et al. Role of spontaneous and sensory orexin network dynamics in rapid locomotion initiation. Prog Neurobiol 187, 101771 (2020).

48. Donegan, D., et al. Hypothalamic Control of Forelimb Motor Adaptation. J Neurosci 42, 6243–6257 (2022).

49. Gonzalez, J.A., et al. Inhibitory Interplay between Orexin Neurons and Eating. Current Biology 26, 2486–2491 (2016).

50. Harris, G.C., Wimmer, M. & Aston-Jones, G. A role for lateral hypothalamic orexin neurons in reward seeking. Nature 437, 556–559 (2005).

51. Hassani, O.K., Krause, M.R., Mainville, L., Cordova, C.A. & Jones, B.E. Orexin Neurons Respond Differentially to Auditory Cues Associated with Appetitive versus Aversive Outcomes. in The Journal of Neuroscience 1747 (Society for Neuroscience, 2016).

52. Zagha, E., et al. The Importance of Accounting for Movement When Relating Neuronal Activity to Sensory and Cognitive Processes. J Neurosci 42, 1375–1382 (2022).

53. James, G., Witten, D., Hastie, T. & Tibshirani, R. An introduction to statistical learning : with applications in R (Springer, New York, 2013).

54. Engelhard, B., et al. Specialized coding of sensory, motor and cognitive variables in VTA dopamine neurons. Nature 570, 509–513 (2019).

55. Musall, S., Kaufman, M.T., Juavinett, A.L., Gluf, S. & Churchland, A.K. Single-trial neural dynamics are dominated by richly varied movements. Nature neuroscience 22, 1677–1686 (2019).

56. Stringer, C., et al. Spontaneous behaviors drive multidimensional, brainwide activity. Science 364, 255 (2019).

57. Wang, A.W., et al. Not everything, not everywhere, not all at once: a study of brain-wide encoding of movement. bioRxiv https://doi.org/10.1101/2023.06.08.544257 (2023).

58. Bimbard, C., et al. Behavioral origin of sound-evoked activity in mouse visual cortex. Nature neuroscience 26, 251–258 (2023).

59. Schultz, W. Dopamine reward prediction-error signalling: a two-component response. Nature reviews. Neuroscience 17, 183–195 (2016).

60. Fiorillo, C.D., Tobler, P.N. & Schultz, W. Discrete coding of reward probability and uncertainty by dopamine neurons. Science 299, 1898–1902 (2003).

61. Schultz, W., Apicella, P. & Ljungberg, T. Responses of monkey dopamine neurons to reward and conditioned stimuli during successive steps of learning a delayed response task. J Neurosci 13, 900–913 (1993).

62. Waelti, P., Dickinson, A. & Schultz, W. Dopamine responses comply with basic assumptions of formal learning theory. Nature 412, 43–48 (2001).

63. Inutsuka, A., et al. The integrative role of orexin/hypocretin neurons in nociceptive perception and analgesic regulation. Sci Rep 6, 29480 (2016).

64. Li, S.B., Nevarez, N., Giardino, W.J. & de Lecea, L. Optical probing of orexin/hypocretin receptor antagonists. Sleep 41 (2018).

65. Legaria, A.A., et al. Fiber photometry in striatum reflects primarily nonsomatic changes in calcium. Nature neuroscience 25, 1124–1128 (2022).

66. Schultz, W. Getting formal with dopamine and reward. Neuron 36, 241–263 (2002).

67. Schultz, W. & Dickinson, A. Neuronal coding of prediction errors. Annu Rev Neurosci 23, 473–500 (2000).

68. Harada, M., Capdevila, L.S., Wilhelm, M., Burdakov, D. & Patriarchi, T. Stimulation of VTA dopamine inputs to LH upregulates orexin neuronal activity in a DRD2-dependent manner. bioRxiv (2023).

69. Kosse, C., Gonzalez, A. & Burdakov, D. Predictive models of glucose control: roles for glucose-sensing neurones. Acta Physiol (Oxf) 231, 7–18 (2015).

70. Goforth, P.B. & Myers, M.G. Roles for Orexin/Hypocretin in the Control of Energy Balance and Metabolism. Curr Top Behav Neurosci 33, 137–156 (2017).

71. Karnani, M. & Burdakov, D. Multiple hypothalamic circuits sense and regulate glucose levels. Am J Physiol Regul Integr Comp Physiol 300, R47–55 (2011).

72. Baimel, C., et al. Orexin/hypocretin role in reward: implications for opioid and other addictions. Br J Pharmacol (2014).

73. Borgland, S.L., et al. Orexin A/hypocretin-1 selectively promotes motivation for positive reinforcers. J Neurosci 29, 11215–11225 (2009).

74. De Lecea, L., et al. Addiction and arousal: Alternative roles of hypothalamic peptides. in Journal of Neuroscience 10372–10375 (Society for Neuroscience, 2006).

75. Mahler, S.V., Smith, R.J. & Aston-Jones, G. Interactions between VTA orexin and glutamate in cue-induced reinstatement of cocaine seeking in rats. Psychopharmacology 226, 687–698 (2013).

76. Burdakov, D. & Gonzalez, J.A. Physiological functions of glucose-inhibited neurones. Acta Physiol (Oxf) 195, 71–78 (2009).

77. Iordanidou, P. & Burdakov, D. Brain glucose feedback predicts food choice (Commentary on Wakabayashi et al.). Eur J Neurosci 43, 1420–1421 (2016).

78. Venner, A., et al. Orexin neurons as conditional glucosensors: paradoxical regulation of sugar sensing by intracellular fuels. J Physiol 589, 5701–5708 (2011).

79. Viskaitis, P., et al. Orexin cells efficiently decode blood glucose dynamics to drive adaptive behavior. bioRxiv (2022).

80. Jordan, R. & Keller, G.B. The locus coeruleus broadcasts prediction errors across the cortex to promote sensorimotor plasticity. Elife 12 (2023).

81. Mickelsen, L.E., et al. Single-cell transcriptomic analysis of the lateral hypothalamic area reveals molecularly distinct populations of inhibitory and excitatory neurons. Nature neuroscience 22, 642–656 (2019).

82. Mickelsen, L.E., et al. Neurochemical Heterogeneity Among Lateral Hypothalamic Hypocretin/Orexin and Melanin-Concentrating Hormone Neurons Identified Through Single-Cell Gene Expression Analysis. eNeuro 4 (2017).

83. Bonnavion, P., Mickelsen, L.E., Fujita, A., de Lecea, L. & Jackson, A.C. Hubs and spokes of the lateral hypothalamus: cell types, circuits and behaviour. in The Journal of Physiology 6443–6462 (Wiley/Blackwell (10.1111), 2016).

84. Soya, S., et al. Orexin modulates behavioral fear expression through the locus coeruleus. Nature communications 8, 1606 (2017).

85. Sakurai, T. The neural circuit of orexin (hypocretin): maintaining sleep and wakefulness. Nature reviews. Neuroscience 8, 171–181 (2007).

86. Tan, Y., et al. Impaired hypocretin/orexin system alters responses to salient stimuli in obese male mice. J Clin Invest 130, 4985–4998 (2020).

87. Horvath, T.L., Diano, S. & van den Pol, A.N. Synaptic interaction between hypocretin (orexin) and neuropeptide Y cells in the rodent and primate hypothalamus: a novel circuit implicated in metabolic and endocrine regulations. J Neurosci 19, 1072–1087 (1999).

88. Horvath, T.L. & Gao, X.B. Input organization and plasticity of hypocretin neurons: possible clues to obesity’s association with insomnia. Cell Metab 1, 279–286 (2005).

89. Rao, Y., et al. Prolonged wakefulness induces experience-dependent synaptic plasticity in mouse hypocretin/orexin neurons. J Clin Invest 117 (2007).

90. Winsky-Sommerer, R., et al. Interaction between the corticotropin-releasing factor system and hypocretins (orexins): a novel circuit mediating stress response. J Neurosci 24, 11439–11448 (2004).

91. De Luca, R., et al. Orexin neurons inhibit sleep to promote arousal. Nature communications 13, 4163 (2022).

92. Adamantidis, A.R., et al. A circuit perspective on narcolepsy. Sleep (2020).

93. Burgess, C.R., Oishi, Y., Mochizuki, T., Peever, J.H. & Scammell, T.E. Amygdala lesions reduce cataplexy in orexin knock-out mice. J Neurosci 33, 9734–9742 (2013).

94. Saper, C.B., Chou, T.C. & Scammell, T.E. The sleep switch: hypothalamic control of sleep and wakefulness. Trends in neurosciences 24, 726–731 (2001).

95. Blomeley, C., Garau, C. & Burdakov, D. Accumbal D2 cells orchestrate innate risk-avoidance according to orexin signals. Nature neuroscience 21, 29–32 (2018).

96. Garau, C., Blomeley, C. & Burdakov, D. Orexin neurons and inhibitory Agrp-->orexin circuits guide spatial exploration in mice. J Physiol 598, 4371–4383 (2020).

97. Bourgin, P., et al. Hypocretin-1 modulates rapid eye movement sleep through activation of locus coeruleus neurons. J Neurosci 20, 7760–7765 (2000).

98. Broberger, C., De Lecea, L., Sutcliffe, J.G. & Hokfelt, T. Hypocretin/orexin- and melanin-concentrating hormone-expressing cells form distinct populations in the rodent lateral hypothalamus: relationship to the neuropeptide Y and agouti gene-related protein systems. J Comp Neurol 402, 460–474 (1998).

99. Peyron, C., et al. Neurons containing hypocretin (orexin) project to multiple neuronal systems. J Neurosci 18, 9996–10015 (1998).

100. Romanov, R.A., et al. Molecular interrogation of hypothalamic organization reveals distinct dopamine neuronal subtypes. Nature neuroscience 20, 176–188 (2017).

101. Wang, Y., et al. EASI-FISH for thick tissue defines lateral hypothalamus spatio-molecular organization. Cell 184, 6361–6377 e6324 (2021).

102. Williams, R.H., Morton, A.J. & Burdakov, D. Paradoxical function of orexin/hypocretin circuits in a mouse model of Huntington’s disease. Neurobiol Dis 42, 438–455 (2011).

103. Bassetti, C.L.A., et al. Narcolepsy - clinical spectrum, aetiopathophysiology, diagnosis and treatment. Nat Rev Neurol 15, 519–539 (2019).

104. Thannickal, T.C., Lai, Y.Y. & Siegel, J.M. Hypocretin (orexin) cell loss in Parkinson’s disease. Brain 130, 1586–1595 (2007).

105. Thannickal, T.C., et al. Reduced number of hypocretin neurons in human narcolepsy. Neuron 27, 469–474 (2000).

106. Petersen, A., et al. Orexin loss in Huntington’s disease. Hum Mol Genet 14, 39–47 (2005).

107. Fronczek, R., et al. Hypocretin (orexin) loss in Alzheimer’s disease. Neurobiol Aging 33, 1642–1650 (2012).

108. Nishino, S., Ripley, B., Overeem, S., Lammers, G.J. & Mignot, E. Hypocretin (orexin) deficiency in human narcolepsy. in Lancet 39-40 (Elsevier B.V., 2000).

109. Kosse, C. & Burdakov, D. Natural hypothalamic circuit dynamics underlying object memorization. Nature communications 10, 2505 (2019).

110. Kim, C.K., et al. Simultaneous fast measurement of circuit dynamics at multiple sites across the mammalian brain. Nat Methods 13, 325–328 (2016).

111. Pachitariu, M., et al. Suite2p: beyond 10,000 neurons with standard two-photon microscopy. https://www.biorxiv.org/content/10.1101/061507v2 (2017).

